# SAMSA2: A standalone metatranscriptome analysis pipeline

**DOI:** 10.1101/195826

**Authors:** Samuel T Westreich, Michelle L Treiber, David A Mills, Ian Korf, Danielle G Lemay

## Abstract

**Background:** Complex microbial communities are an area of rapid growth in biology. Metatranscriptomics allows one to investigate the gene activity in an environmental sample via high-throughput sequencing. Metatranscriptomic experiments are computationally intensive because the experiments generate a large volume of sequence data and the sequences must be compared with many references.

**Results:** Here we present SAMSA2, an upgrade to the original Simple Annotation of Metatranscriptomes by Sequence Analysis (SAMSA) pipeline that has been redesigned for use on a supercomputing cluster. SAMSA2 is faster due to the use of the DIAMOND aligner, and more flexible and reproducible because it uses local databases. SAMSA2 is available with detailed documentation, and example input and output files along with examples of master scripts for full pipeline execution.

**Conclusions:** Using publicly available example data, we demonstrate that SAMSA2 is a rapid and efficient metatranscriptome pipeline for analyzing large paired-end RNA-seq datasets in a supercomputing cluster environment. SAMSA2 provides simplified output that can be examined directly or used for further analyses, and its reference databases may be upgraded, altered or customized to fit the specifics of any experiment.

## 1. Background

High-throughput sequencing methods are now used to identify both culturable and unculturable microbial species. Although 16S ribosomal profiling is still most commonly used, there is an increasing shift towards using more comprehensive sequencing methods such as metagenomics and metatranscriptomics. Metagenomics—sequencing of all DNA from a diverse sample—reveals which microbes are present. Metatranscriptomics–sequencing of all RNA from a diverse sample captures all gene expression, giving a view of which microbes are active and what they are doing. Despite the power of metatranscriptomics, there are still relatively few bioinformatics tools designed to handle this complex type of data.

Simple Annotation of Metatranscriptomes by Sequence Analysis (SAMSA) was the first open-source bioinformatics pipeline designed specifically for metatranscriptomic data [1]. The first version of SAMSA was built specifically for researchers with minimal bioinformatics experience who may not have a supercomputing cluster available for their use. SAMSA worked in conjunction with MG-RAST [2], a public annotation service capable of handling metagenomic or metatranscriptomic data. SAMSA improved on MG-RAST by offering preprocessing and paired-end merging of RNA sequences, uploading to MG-RAST, downloading annotations, and analyzing the results. One downside of MG-RAST is that the software and database are not under control of the user. For maximum reproducibility, one must have complete control over all parameters, and this is not possible with a web service.

SAMSA2 is designed for cluster computing. The software and databases can be completely containerized for reproducibility. SAMSA2 uses DIAMOND [3] for aligning sequences, which greatly increases its speed. SAMSA2 handles end-to-end analysis of metatranscriptomes from quality control of sequencing reads through publication-ready images. SAMSA2 is packaged with documentation and sample data.

## 2. Implementation

### 2.1. Recommended sequencing parameters

To provide an accurate view of all activity occurring within the complex and varied environment of a microbiome, it is important to obtain sufficient depth of sequencing, along with reads of length necessary to avoid erroneous alignments. Longer reads are generally more advantageous, as they are less likely to be erroneously aligned to references. For all tests of SAMSA, paired-end 100bp reads were obtained using an Illumina HiSeq 4000 (Illumina, Inc., San Diego, CA). The SAMSA pipeline includes a merging step to combine overlapping pairedend reads; while this step can be skipped if using single-end samples, the shorter read lengths can contribute to incorrect annotations, if they are not at least 150bp.

In previous work [1], we established that between 10 and 20 million annotations—which translates to 40–50 million raw (unannotated) reads in stool metatranscriptomes—ensures best results when estimating the abundance of transcripts and obtaining stable measures of activity. These minimum annotation counts likely depend significantly on the stability of the chosen microbiome, as well as on the magnitude of the target signal or reaction. For tests run on SAMSA2, starting files containing 50–60 million reads were used; these files were obtained by running four samples per lane on the Illumina HiSeq 4000.

Given the high abundance of ribosomal RNAs (rRNAs) naturally present in total RNA extracted from microbiome samples, ribodepletion is strongly encouraged. A digital ribodepletion step is included in the SAMSA pipeline, but physical ribodepletion before sequencing ensures that a higher number of mRNAs are sequenced. Ribodepletion kits do not effectively remove all rRNA from samples, but they greatly reduce (>80%) the number of sequenced rRNA reads that must later be discarded from bioinformatic analysis. In summary, it is recommended that metatranscriptomes be ribodepleted and sequenced in a paired-end format of at least 100bp with a minimum of 40 million reads per sample.

### 2.2. Preprocessing

An overview of the flow of data through the SAMSA version 2.0 pipeline is in Figure 1. The first step in the SAMSA pipeline is to merge paired-end files, if this sequencing type was used. PEAR is a fast and efficient paired-read merger [4] that can be installed as a precompiled binary. When used on two paired-end files, it creates a merged output containing all reads with overlap, as well as two notCombined files that contain forward and reverse unassembled reads, respectively. Generally, only the merged reads are used in the remainder of the pipeline, although if a high number of reads cannot be merged, it may be advisable to use both the merged and the forward notCombined read sets to ensure that an adequate total number of annotations is obtained.

**Figure 1.**
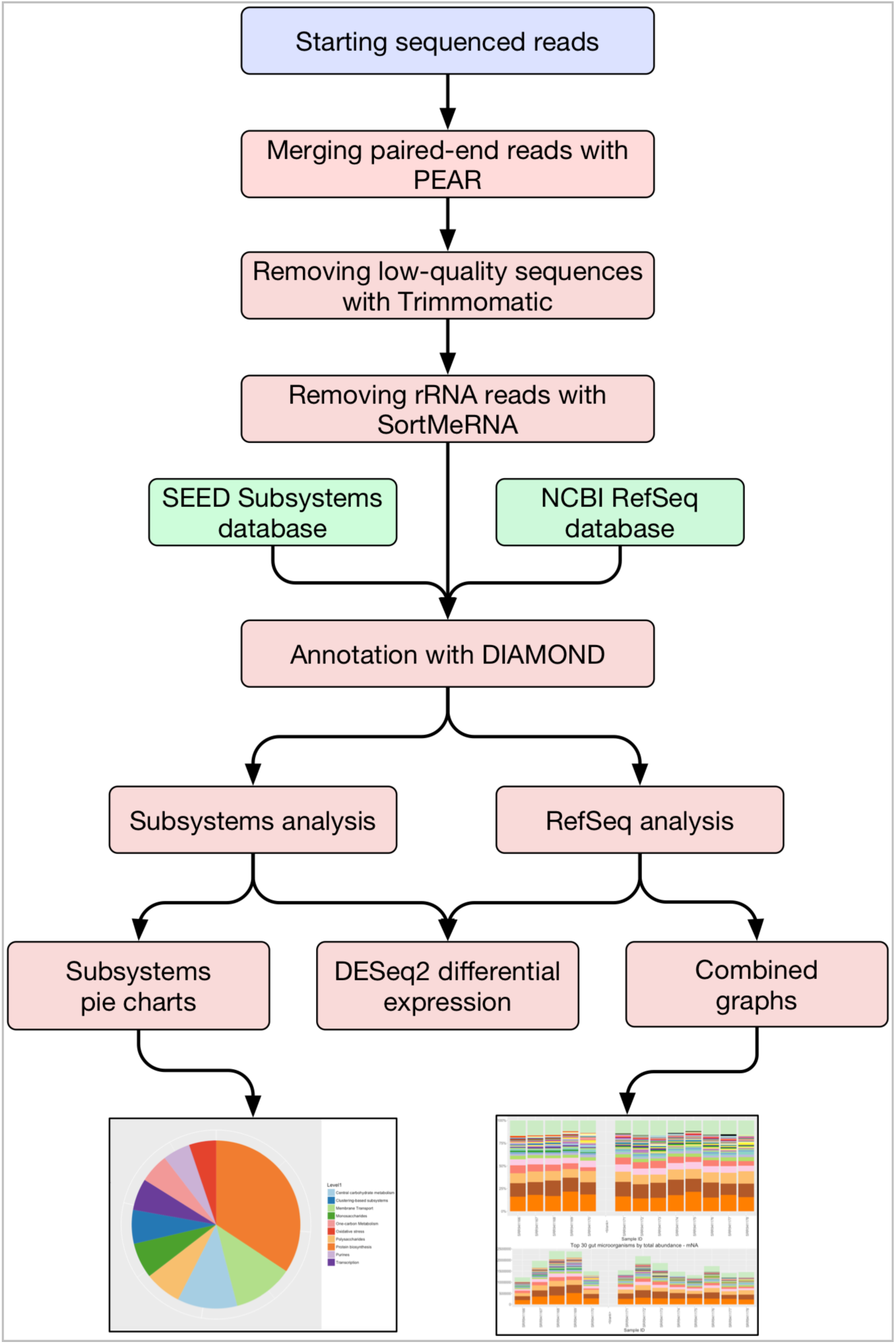
SAMSA2 metatranscriptome analysis pipeline.

Following paired-end read merging (if necessary), the files must be cleaned to remove low-quality sequences and/or adaptor contamination. Trimmomatic [5] is a flexible read trimmer designed for Illumina sequence data, which can both remove low-quality sequences, and identify and remove any potential adaptor contamination. Trimmomatic produces a cleaned output file that is ready for the next step in preprocessing.

Finally, although the majority of ribosomal sequences are hopefully removed from the metatranscriptome before sequencing, SortMeRNA [6] provides another digital sweep, ensuring that ribosomal reads are removed before annotation. Removal of ribosomal reads at this step reduces the total number of sequences that will be annotated, increasing downstream speed and reducing file sizes. Another reason to discard these sequences is that the remaining ribosomal reads cannot be used to assess organism activity because the ribodepletion is biased towards certain organisms [1]. SortMeRNA installs with several reference rRNA databases for both bacterial and eukaryotic ribosome sequences. SortMeRNA outputs both identified ribosomal sequences and sequences that do not match against ribosomes; these “other” sequences are carried forward to the annotation step in SAMSA.

### 2.3. Annotation

SAMSA2 uses DIAMOND [3], a superfast BLAST-like algorithm, to perform annotations against one or more selected reference databases. DIAMOND is designed specifically for annotating large numbers of read sequences against a reference simultaneously, with speeds up to 10,000 times faster than traditional BLAST.

Before input sequences can be annotated by DIAMOND, however, reference databases must be created. One advantage of SAMSA2 is that any set of translated sequences in FASTA format may be provided to DIAMOND and converted into a reference database. Instructions for creating DIAMOND-searchable databases are included in the documentation provided with SAMSA2, along with links and more detailed installation instructions for creating databases from NCBI’s RefSeq database [7] and the SEED Subsystems hierarchical database [8] for functional activities. It is also possible to add any other database, such as the Carbohydrate Active Enzyme (CAZy) database [9], available for download in a translated FASTA format.

Once a database has been created and converted into a DIAMOND-searchable binary format, experimental sequence files can be annotated against this reference. DIAMOND runs in two steps; the first step creates a binary output containing all matches for input reads against the reference database, while the second step converts the binary outfile to a viewable BLAST m8 data table. This m8 formatted data file is used in the downstream steps.

### 2.4. Aggregation and downstream processing

Annotated files returned from DIAMOND searches are returned in plaintext; each sequence from the infile that was annotated to a sequence in the reference database occupies one line in the outfile. The next step is to aggregate these large, line-by-line files into a condensed, sorted summary of output.

Using Python, SAMSA2 converts the DIAMOND output into a sorted abundance count, creating two files—one file is for organism annotations, whereas the other file contains functional annotations of the same reads. When reads are annotated against a hierarchical database such as SEED Subsystems, this sorted abundance count also includes all levels of hierarchy information.

These sorted abundance counts are saved as the final step of the automated workflow for the SAMSA2 pipeline. These count files can be viewed directly to identify most active organisms or functions within a metatranscriptome and are also used by R in the final step of the SAMSA2 pipeline for statistical analysis.

The final step in the SAMSA2 pipeline is analysis using R scripting for calculating differential expression and generating tables and graphs, treating the sorted abundance counts from the aggregation step of the SAMSA2 pipeline as input. Calculation of significant differences at both the organism and the function level for each database is performed automatically in the batch submission script, whereas the outputs may be imported into an interactive R session for further analysis or figure generation.

Multiple R scripts are included with the SAMSA2 pipeline, each designed to carry out a single analysis action on the processed data. These included R scripts can perform the following actions: determine statistically significant differentially active organisms or functions between two groups of samples (such as an experimental condition compared with normal controls), calculate microbial diversity statistics for each metatranscriptome, create heatmaps of organism or functional activity, create PCA plots based on organism or functional activity, and create stacked bar graphs for each metatranscriptome, displaying the most abundant functions or organisms within each metatranscriptome (Figure 2). For metatranscriptomes annotated against the Subsystems hierarchical database, included R scripts can identify statistically significant differentially expressed categories at each hierarchy level, and can create pie charts for each metatranscriptome to show relative activity matching each functional activity category (Figure 3).

**Figure 2.**
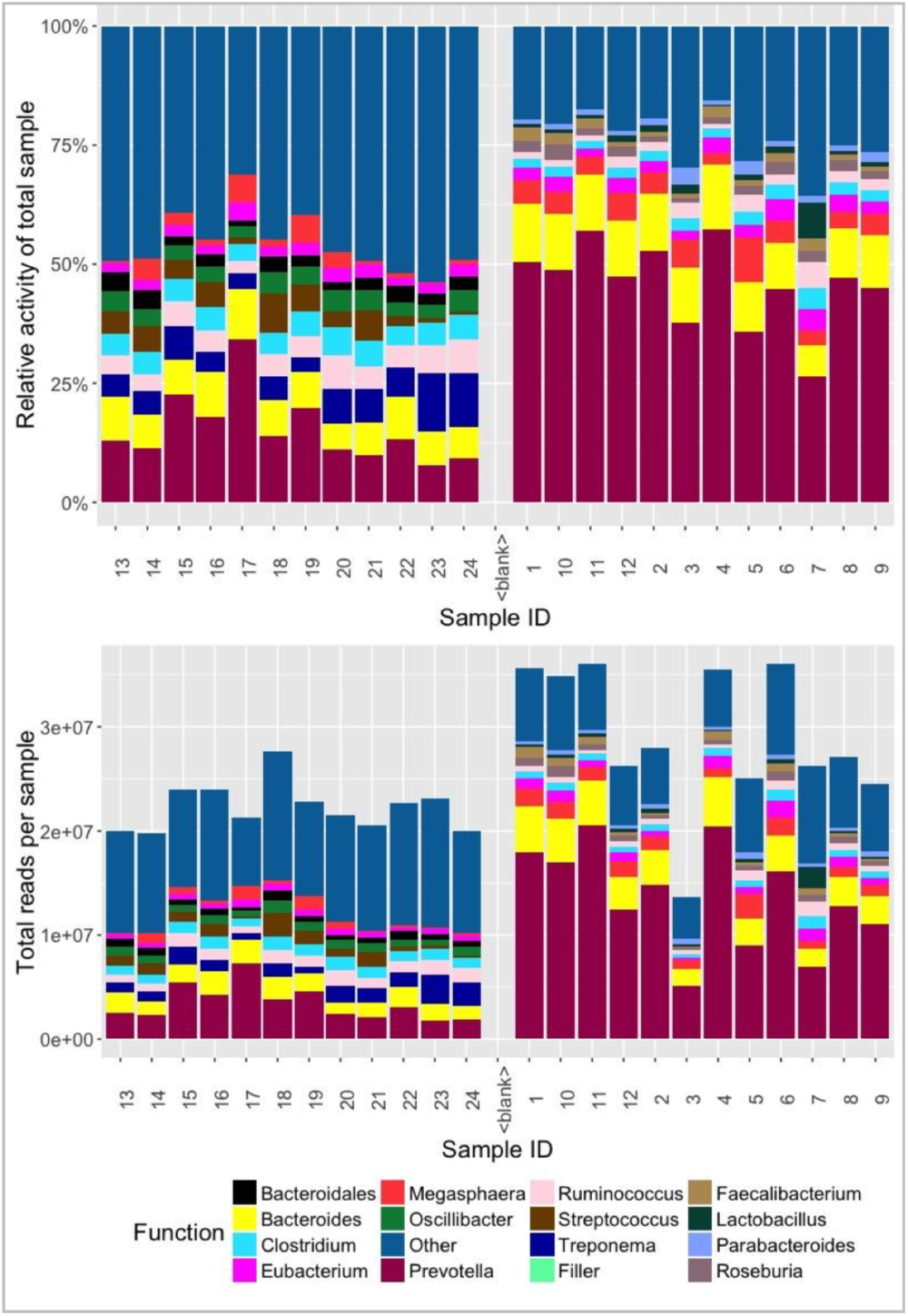
Sample organism output from SAMSA2 results. R scripts for generating stacked bar graphs, heatmaps, PCA plots, and other visuals are included in the SAMSA package.

**Figure 3.**
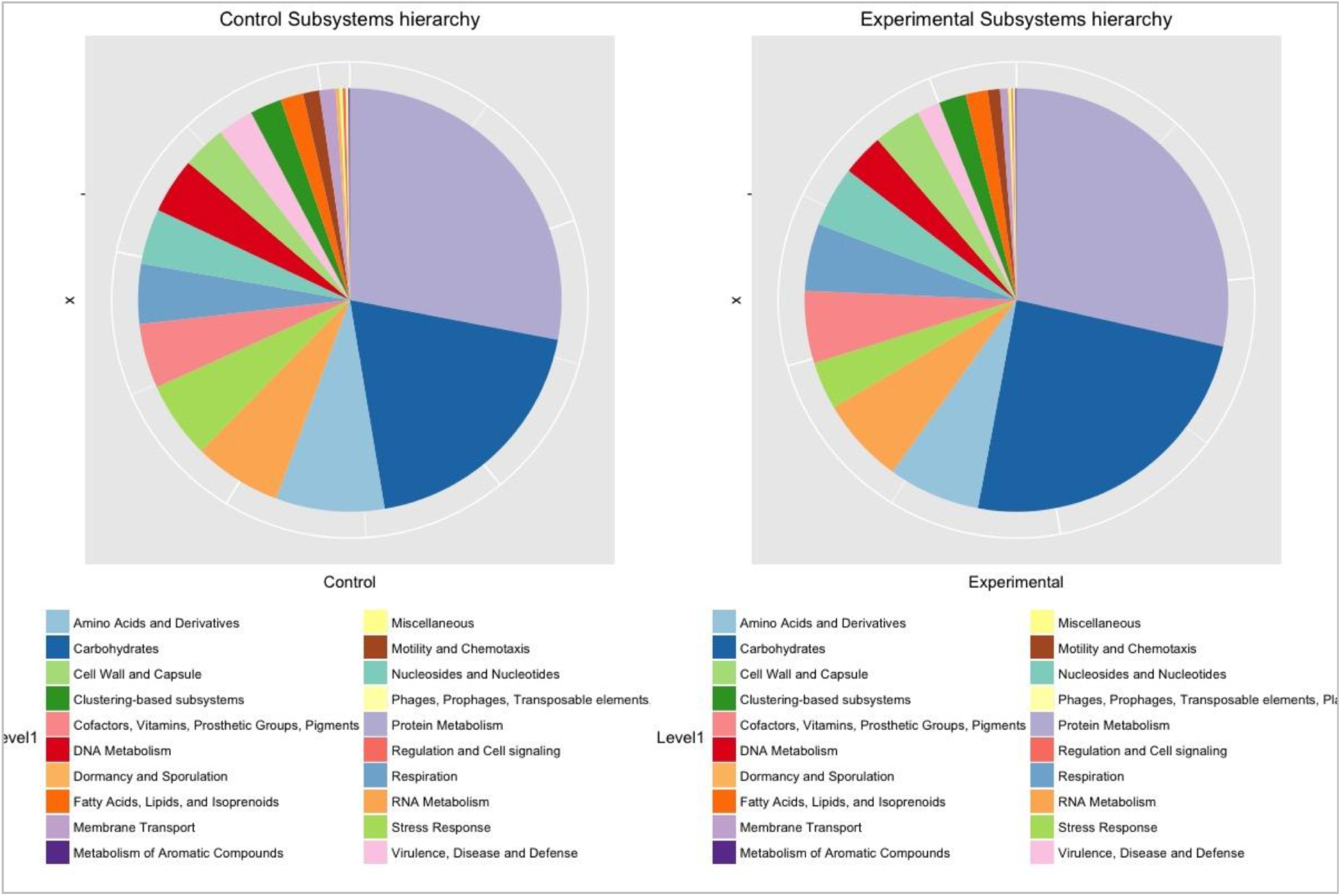
SEED Subsystems annotation pie charts at hierarchy level 1.

## 3. Results

### 3.1 Improved speed and accuracy for metatranscriptome analysis

Although the earlier version (1.0) of SAMSA provided a complete metatranscriptome analysis pipeline, it depended on a public web service (MG-RAST) for the annotation step, which was a major roadblock with respect to speed due the growing popularity of this resource. In contrast, SAMSA2 is standalone in the sense that the tool dependencies are downloadable and can be run on the user’s own compute resources. SAMSA2’s annotation speed was tested with various file sizes. SAMSA2 scales relatively linearly as metatranscriptome size increases (Figure 4A).

**Figure 4.**
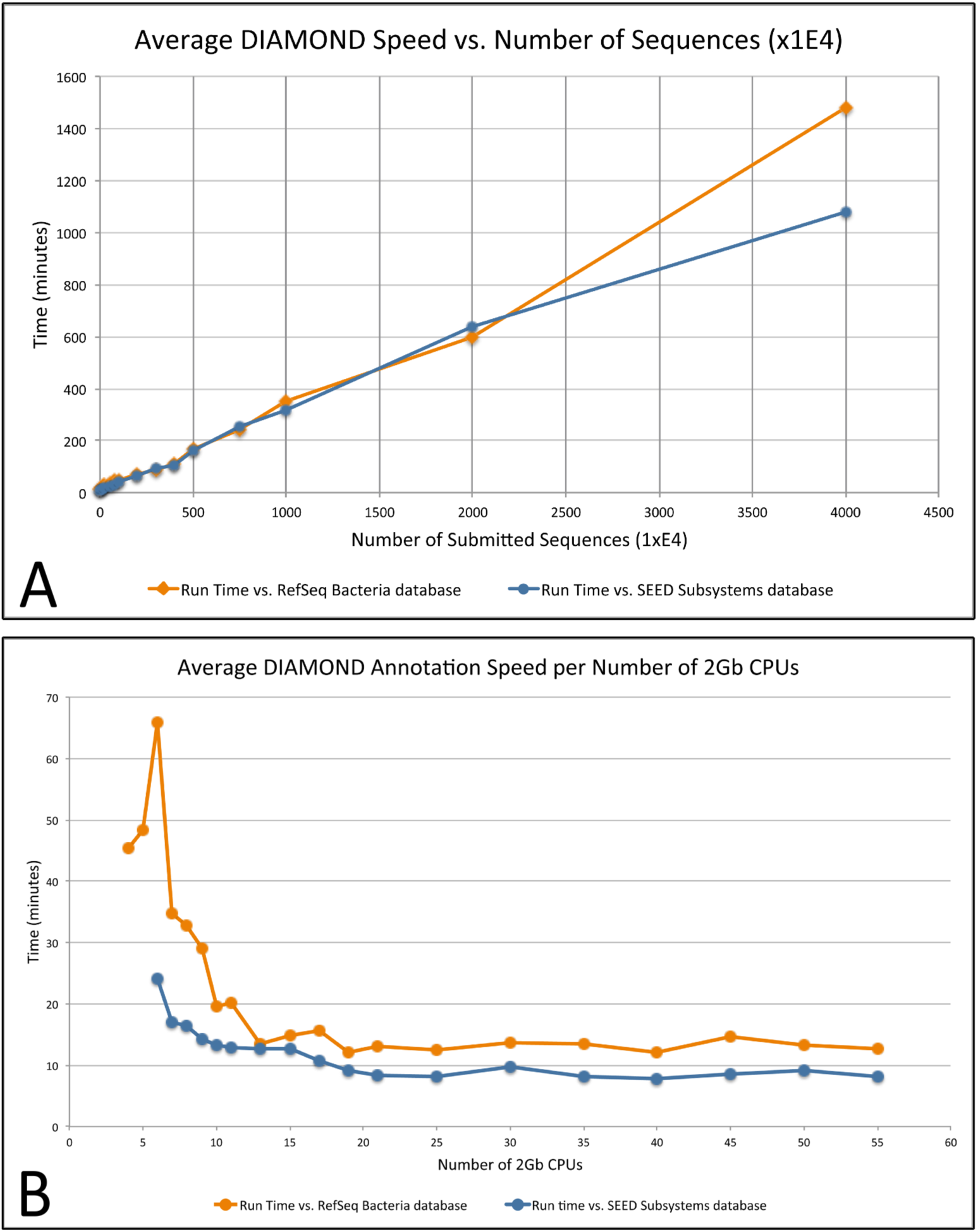
**(A)** DIAMOND annotation time increases linearly as more input sequences are added, allowing the estimation of total annotation time. For this test, all files ran with 30 CPUs, each with 2 GB RAM. **(B)** Annotation speed relative to allocated memory: Higher RAM allocation allows DIAMOND to hold more of the reference databases in memory, speeding up pipeline annotation up to the point where the database is fully in memory; all files in this test contained 50,000 sequences each.

SAMSA2 was tested with increasing numbers of CPU and increasing RAM size. These experiments suggest that a minimum of 10 CPUs/20 GB of RAM are needed to efficiently process even small sequence files against NCBI’s non-redundant Bacteria database,approximately 6 GB (see Figure 4B). If a larger reference database is used, even more RAM will be needed.

Additionally, SAMSA2 yields consistently high accuracy scores. Using simulated metatranscriptomes made from reference reads through the use of MetaSim [10], the accuracy of SAMSA2’s matching was determined for both organism and function. Organism matching was evaluated by comparing SAMSA2’s predicted organism to the actual organism from which the starting sequence was obtained, whereas function matching was evaluated by comparing the predictive function of the entire gene sequence to the functions predicted from the smaller read sequences used as input for SAMSA2 testing. Overall, SAMSA2 had high accuracy for sequences from a selection of the most common organisms identified in gut microbiome samples, with approximately an 87.5% accurate prediction of organism at the species level. Selecting at the genus level, SAMSA2 yielded approximately a 95% accuracy at the annotation step.

Examining annotation accuracy of functions is difficult, given the lack of standard naming conventions for functional activity of genes. To overcome this challenge, functional annotations were performed against the SEED Subsystems database, which offers the unique advantage of providing a hierarchy for each function, with each specific functional annotation grouped under several more general categories. Functional annotations of reads generated by MetaSim from supplied starting sequences were annotated and then compared against the SEED Subsystems annotations of full sequences. Although there are often difficulties with predicting the exact function from partial reads, as expected (approximately 76% accuracy), the SAMSA2 pipeline overall shows excellent accuracy for predicting hierarchy at all higher levels of organization (greater than 95% accuracy). Given that other sequencing approaches generally fail to provide any functional information about the microbiome, SAMSA2’s impressive accuracy at predicting both organism and function for each read makes it powerful for providing a complete picture of a microbiome community.

### 3.2. Ability to customize index database

One challenge faced by nearly all bioinformatics pipelines is that some databases are not public and are restricted by institutional paywalls, whereas others may contain suspect annotations that could prove to be erroneous. Compounding this issue is the differing rate of updates to various databases; the release version of NCBI’s RefSeq database [7] used by one pipeline may be different from the version used by another similar tool, yielding differing annotation results even from the same sequencing reads as input.

Version 1 of SAMSA relied upon the databases compiled by MG-RAST. MG-RAST uses internal identifiers to combine multiple databases, including NCBI’s RefSeq, UniProt, SEED Subsystems, KEGG Orthologs, GreenGenes, SILVA SSU, and others [7, 8, 11-14]. A single internal identifier mapped to the corresponding entry for a read in each of these databases allows cross-database comparisons. Although this approach is an excellent method to compensate for a lack of a “global thesaurus” between different databases, it does not solve issues with version compatibility. MG-RAST’s version of KEGG Orthologs, for example, is the last available version before KEGG made its database private in 2011, and is thus likely out of date [15].

SAMSA2 provides a solution to the database issue by offering the ability to add or even create, custom databases that can be indexed and searched against for metatranscriptome annotation. Any database that can be downloaded in or converted to FASTA protein format, can be incorporated into SAMSA2 and used as a reference database. Downstream scripting is currently optimized to run using NCBI’s RefSeq database for organism and functional annotation, with the option to include the publicly available SEED Subsystems database [8] for hierarchical clustering of functional annotations. It is also possible to add any other available database, including more specific databases such as the Carbohydrate Active Enzyme (CAZy) database [9].

To add a custom database to the SAMSA2 pipeline, the database must be available as an unencrypted text file in FASTA format, where the header contains an internal ID, the function name, and the organism name in brackets. A single command converts this file into a DIAMOND-indexed encrypted database file, which can then be used for analysis of metatranscriptome files. Because these databases are stored on the cluster or server running SAMSA2, there is no limit to how many index databases or versions of databases can be used or consulted.

### 3.3. Functional annotations and organism annotations for each input read

An important feature of metatranscriptome analysis is the ability to trace each read through the pipeline, determining exactly which sequences are assigned to a specific annotation. This tracking allows for more powerful analyses, such as determining all functions performed by a specific organism or clade, determining which organisms are carrying out a specific function or category of functions or determining the ability to evaluate for potentially differing annotations from different databases.

SAMSA2 generates outfiles after each step in the pipeline, ensuring that version history and each entry read are tracked. The outfile from the annotation step contains each read’s original identifier assigned to it by the sequencing machine, as well as its organism and function annotations. Although these outfiles are later aggregated to create sorted abundance counts, additional scripting allows SAMSA2 users to alter the selection parameters for the creation of these sorted abundance counts files, specifying whether they want to include either all annotations or a subset (selecting either by organism or by functional keyword or category for sorting and selection).

By minimizing loss of information, SAMSA2 allows users to perform the longest and most computationally expensive step of annotation only once per database, after which they may return as many times as needed to the outfiles for additional analyses. There is no need to rerun the entire sample files if changes need to be made or additional reads are to be added; those can simply be aggregated on to existing results to save time and computational resources.

### 3.4. Sorting of functions into hierarchical categories using SEED Subsystems

The ability to annotate mRNA sequence data from a microbial community not just to organism, but also to its function, is incredibly powerful. However, in such a big data context, a traditional approach, such as the results returned by a BLAST search, quickly proves overwhelming. With a single metatranscriptome likely to return anywhere from tens to hundreds of thousands of unique functions when mapped against a traditional index database such as RefSeq, determining the general activities occurring within this environment becomes next to impossible.

To overcome this issue, SAMSA2 incorporates the open-source SEED Subsystems database [8], which is uniquely organized through hierarchical classification. Each sequence in the SEED Subsystems database has a unique functional annotation but is also grouped into increasingly broad categories at four levels of hierarchy. At the highest hierarchy level, a single category may include tens of thousands of different specific functions.

Similar to other reference databases, the SEED Subsystems database may be downloaded and indexed by DIAMOND for use in annotation. Annotation against the SEED Subsystems database provides the exact functional match within the Subsystems database, but additional scripting allows the Subsystem functional hierarchy to be retrieved for each annotation. This allows for analysis of the metatranscriptome at each level of hierarchy, including determination of significant differential expression and graphical representation of relative functional activity at any chosen hierarchy level (see Figure 3). While the change in many specific functions may be too small to be noted as significantly differentially expressed between experimental conditions, the broader category of functional activity reveals the larger-scale shift in activity within the community. Full instructions are included in the SAMSA2 documentation for downloading, compiling, and using the SEED Subsystems database for examining functional hierarchies of annotations.

### 3.5. Subdividing metatranscriptome data to obtain functional activity by specific organism

An important factor for microbiome analysis is the ability to perform a joint analysis of both organism and function. Without knowing which organisms are performing specific activities, any examination of microbiome activities is greatly limited. Metatranscriptomics offers the ability to determine for reads matching a specific functional activity or category, which organisms produced those sequences. SAMSA2 provides this information by offering optional scripts to segment the data by organism or by function or functional category. Results from an entire microbiome may be subdivided to examine only the activities of reads annotated to a single genus or species, and functional annotations in a specific group may be examined to determine which organisms produce those transcripts. By matching the read IDs assigned to transcripts against the reference databases, which contain both organism and functional annotations, it is possible to obtain both the organism and functional results for each specific read in the entire metatranscriptome.

## 4. Discussion

Study of complex microbial environments, which may contain many different interacting organisms, requires large amounts of data to fully understand. Current approaches, such as 16S rRNA sequencing, can provide a broad overview of which groups of microbes are present in an environment, but they fail to offer enough resolution to differentiate between closely related genera or species, and they provide no information about potential activities being performed by members of the microbiome; further verification is required to answer questions of function, usually through labor-intensive microbial phenotyping studies.

Although metatranscriptomics generally requires higher initial sample quality and has higher costs in sequencing and processing, it offers a full and unparalleled view of a complex microbial environment, offering detailed insights into both the organisms present and their current transcriptional activity. However, like many new and emerging advances in scientific analysis, metatranscriptomics is often limited by the lack of available pipelines and bioinformatics tools for processing these complex and extremely large datasets. The first metatranscriptomics analysis papers described workflows but did not provide code or a software program [16, 17]. The first published open-source pipeline, SAMSA [1], utilized the MG-RAST server [2] for the mapping step, enabling users to analyze metatranscriptomes without access to compute resources. Similarly, a web-based tool called COMAN enables users to analyze metatranscriptomes without access to compute resources [18]. Although SAMSA offers superior functional annotation, via the SEED database [8] that is part of MG-RAST [2], both SAMSA and COMAN have similar drawbacks in that they are dependent upon a web service that may become oversubscribed and slow and they fail to support mapping reads to custom reference databases [18]. Another recent tool that is not a web-service, MetaTrans [19], relies on rRNA for organism identification, necessitating a lack of biological ribodepletion and thus reducing the level of captured functional data from mRNA. Our previous work demonstrated that approximately 50 million paired-end reads per sample are needed to accurately quantify transcripts in stool samples [1] when the total RNA has been ribodepleted. For un-ribodepleted samples, the majority of RNA will be rRNA, so the sample would require hundreds of millions of reads for accurate functional quantification. We also previously demonstrated that ribodepleted RNA has biases, so the rRNA must be discarded rather than used for quantification [1]. Finally, some recent metatranscriptome pipelines (anvi’o, [20], IMP, [21]) rely on BLAST [22] for annotations, which may be unacceptably slow when processing multiple millions of sequences per sample. DIAMOND [3], which is used in SAMSA2, is 10,000 times faster than BLAST. Therefore, while there are several tools available today to analyze metatranscriptomes, none offers SAMSA2’s ability to be independently deployed on a high-performance compute cluster or workstation and to use custom databases.

Version 1.0 of SAMSA was intended for use by life science researchers who had limited command line and coding experience, providing straightforward, completely open source tools for the analysis of metatranscriptomic datasets. Many researchers don’t have access to a local computing cluster or a remote server; SAMSA 1.0 allows these researchers to analyze the large metatranscriptome data files by outsourcing the heavy lifting of annotation to the external MG-RAST server. Downstream SAMSA 1.0 programs condense and reduce this huge amount of information and can run on a personal computer.

However, it is becoming easier and cheaper to gain access to bioinformatics supercomputing resources, such as through Amazon Web Services (AWS) or institutional compute clusters. Researchers who have access to these resources need a metatranscriptome analysis solution that is faster and more customizable than that provided by SAMSA 1.0. For this reason, we created the updated SAMSA2; the pipeline still uses the same steps for analyzing metatranscriptomes—preprocessing, annotation, aggregation, and analysis. By switching to local programs that take advantage of the increased processing power of a cluster or supercomputing instance, SAMSA2 is able to offer faster analysis of metatranscriptome files, while providing even more options for in-depth examination than were previously available.

SAMSA2’s speed and customizability gives it advantages over other pipelines currently available for metatranscriptome analysis. Other pipelines still rely on ribosomal sequences for identifying organisms within the microbiome, which leads to either massively over-sequencing the ribosomal profile, failing to obtain enough mRNA reads [19], or lacking the ability to customize the source databases used as references [23]. Additionally, the ability to add custom databases that may be indexed and used as references allows SAMSA2 to be upgraded when newer database versions become available without having to reinstall the entire pipeline and associated programs, and offers version control so that previous analyses may be repeated without concern over reference database version.

As researchers continue to look deeper at microbiomes, seeking to obtain both greater specificity in identifying the organisms active in the environment, and activities and functions of these organisms without needing extensive additional metabolomics analysis, metatranscriptomes will continue to grow to be a more popular option. The costs of sequencing large numbers of transcripts and of digital bioinformatics processing are both decreasing rapidly as sequencer technologies and innovations in computer design combine to reduce prices. As these costs drop, one of the largest challenges that will face microbiome researchers will be identifying flexible and adaptable bioinformatics pipelines for data analysis that do not require extensive programming experience or costly subscriptions to use. SAMSA2 aims to be one potential pipeline option with the flexibility to adapt to the needs and resources of a wide range of biological investigations.

## 5. Conclusion

We have developed SAMSA2, a complete metatranscriptome analysis pipeline that allows for rapid and customizable annotation and analysis of metatranscriptome (RNA-seq for complex microbial communities) data. SAMSA2 can be installed on a local or cloud server and supports incorporation of multiple reference databases, including NCBI’s RefSeq for organism and specific function identification, and SEED Subsystems for hierarchical ontologies of functional activity groups. SAMSA2 is open source (https://github.com/transcript/samsa2).

## Availability and requirements

### Project name: SAMSA2

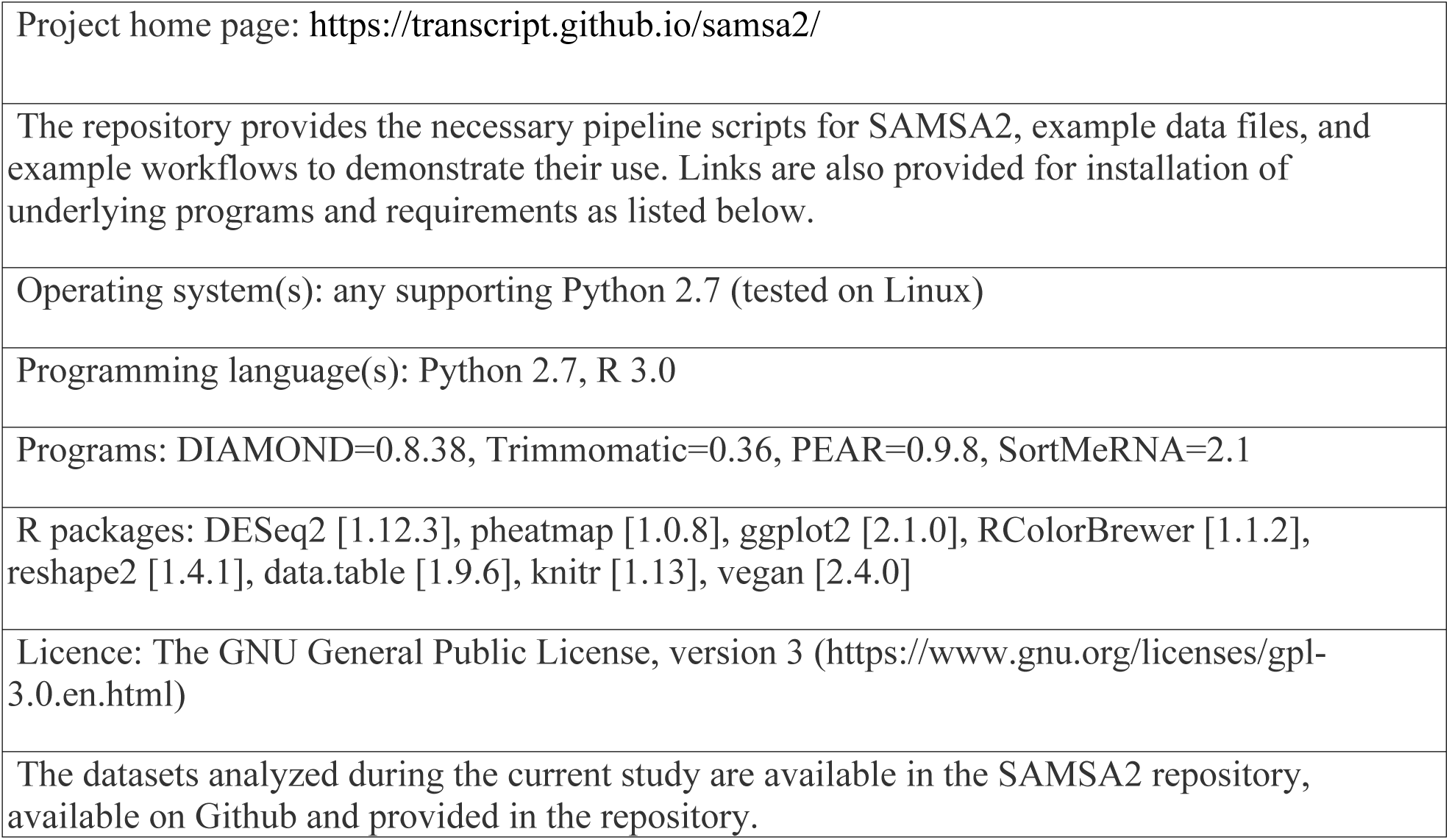

|  |
| --- |
| Project home page: https://transcript.github.io/samsa2/ |
| The repository provides the necessary pipeline scripts for SAMSA2, example data files, and example workflows to demonstrate their use. Links are also provided for installation of underlying programs and requirements as listed below. |
| Operating system(s): any supporting Python 2.7 (tested on Linux) |
| Programming language(s): Python 2.7, R 3.0 |
| Programs: DIAMOND=0.8.38, Trimmomatic=0.36, PEAR=0.9.8, SortMeRNA=2.1 |
| R packages: DESeq2 [1.12.3], pheatmap [1.0.8], ggplot2 [2.1.0], RColorBrewer [1.1.2], reshape2 [1.4.1], data.table [1.9.6], knitr [1.13], vegan [2.4.0] |
| Licence: The GNU General Public License, version 3 (https://www.gnu.org/licenses/gpl-3.0.en.html) |
| The datasets analyzed during the current study are available in the SAMSA2 repository, available on Github and provided in the repository. |

## Abbreviations

- SAMSA, Simple Analysis of Metatranscriptomes through Sequence Annotation, the documented pipeline.
- mRNA, messenger RNA, single-stranded genetic material that codes for the creation of proteins.
- MG-RAST, the MetaGenomics Rapid Annotation using Subsystems Technology server, a public analysis pipeline for handling metagenome and metatranscriptome datasets.
- PCA, principal component analysis.
- DIAMOND, a BLAST-like sequence aligner that uses translated sequence queries to greatly increase annotation speed over traditional BLAST.
- SEED, a protein database that seeks to group sequences into hierarchical categories, created by the Fellowship for Interpretation of Genomes (FIG group).

## Declarations

### Availability of data and material

All supporting data and code, as well as full access to the source code for SAMSA2, is freely available through Github and may be reused under the Creative Commons Attribution 4.0 International License (http://creativecommons.org/licenses/by/4.0/), which permits unrestricted use, distribution, and reproduction in any medium, provided you give appropriate credit to the original author(s) and the source, provide a link to the Creative Commons license, and indicate if changes were made. The Creative Commons Public Domain Dedication waiver (http://creativecommons.org/publicdomain/zero/1.0/) applies to the data made available in this article, unless otherwise stated.

## Authors’ contributions

Pipeline designed and built by STW, based on previous work IK and DGL reviewed code. STW, MLT, and DGL drafted the manuscript. MLT tested code. STW, DGL, DAM, MLT, and IK interpreted data and contributed to the manuscript. All authors read and approved the final manuscript.

## Acknowledgements

The authors thank the UC Davis Bioinformatics Core for technical assistance

The authors also thank Cora Morgan for editorial assistance on preparing the manuscript.

